# Discrete GPCR-triggered endocytic modes enable β-arrestins to flexibly regulate cell signaling

**DOI:** 10.1101/2022.07.13.499995

**Authors:** Benjamin Barsi-Rhyne, Aashish Manglik, Mark von Zastrow

## Abstract

β-arrestins are master regulators of cellular signaling that operate by desensitizing ligand-activated G protein-coupled receptors (GPCRs) at the plasma membrane and promoting their subsequent endocytosis. The endocytic activity of β-arrestins is ligand-dependent, triggered by GPCR binding, and increasingly recognized to have a multitude of downstream signaling and trafficking consequences that are specifically programmed by the bound GPCR. However, only one biochemical ‘mode’ for GPCR-mediated triggering of the endocytic activity is presently known– displacement of the β-arrestin C-terminus (CT) to expose CCP-binding determinants that are masked in the inactive state. Here we revise this view by uncovering a second mode of GPCR-triggered endocytic activity that is independent of the β-arrestin CT and, instead, requires the cytosolic base of the β-arrestin C-lobe (CLB). We further show each of the discrete endocytic modes is triggered in a receptor-specific manner, with GPCRs that bind β-arrestin transiently (‘class A’) primarily triggering the CLB-dependent mode and GPCRs that bind more stably (‘class B’) triggering both the CT and CLB-dependent modes in combination. Moreover, we show that each mode has opposing effects on the net signaling output of receptors– with the CLB-dependent mode promoting rapid signal desensitization and the CT-dependent mode enabling prolonged signaling. Together, these results fundamentally revise understanding of how β-arrestins operate as efficient endocytic adaptors while facilitating diversity and flexibility in the control of cell signaling.

## Introduction

β-arrestins were named for their ability to desensitize signaling by binding to ligand-activated GPCRs and physically blocking heterotrimeric G protein coupling (Lohse et al. 1990), a function similar to the ‘arrest’ of signaling mediated by binding of visual arrestin to the light-activated GPCR rhodopsin (Kühn and Wilden 1987). β-arrestin, unlike visual arrestin, has the additional ability to act as an endocytic adaptor protein by associating with clathrin-coated pits (CCPs) after binding to a GPCR (Goodman et al. 1996; Ferguson et al. 1996). This association promotes clustering of GPCR/β-arrestin complexes on the plasma membrane leading to the subsequent internalization of complexes via clathrin-mediated endocytosis. The prevailing current view is that all GPCRs trigger β-arrestins’ endocytic activity in the same way, by displacing the β-arrestin C-terminus (CT) to unmask endocytic determinants contained therein (Extended Data Fig. 1a) (Xiao et al. 2004; Nobles et al. 2007; Milano et al. 2002; Shukla et al. 2013).

Compelling experimental support for this mode of regulated endocytosis has accumulated over the past 25 years, including the identification of specific clathrin (CHC) and AP2 (AP2β)-binding elements present in the CT which promote the clustering of GPCR/β-arrestin complexes in CCPs (Goodman et al. 1996; Laporte et al. 2000; Orsini and Benovic 1998; Krupnick et al. 1997; Ferguson et al. 1996; Kim and Benovic 2002; Burtey et al. 2007; Ford, Schmid, and McMahon 2007; Kang et al. 2009). However, for most of this time, β-arrestins were thought to regulate signaling only from the cell surface. It is now clear that β-arrestins are more flexible regulators and transducers, and that they have the capacity to promote as well as attenuate signaling from the plasma membrane and endomembranes (Calebiro et al. 2009; Ferrandon et al. 2009; Feinstein et al. 2013; Irannejad et al. 2013; DeFea et al. 2000; McDonald et al. 2000; Terrillon and Bouvier 2004). Furthermore, it is now also widely recognized that β-arrestins produce different downstream signaling effects depending on GPCR-specific differences in the composition, stability and/or conformation of the complex that they form with GPCRs (Lee et al. 2016; Latorraca et al. 2020; Mayer et al. 2019; Nobles et al. 2011; Nuber et al. 2016; Yang et al. 2015, 2018; Asher et al. 2022). In light of the high level of diversity and flexibility that is presently recognized to exist at the GPCR/β-arrestin interface, we wondered if release of the β-arrestin CT is sufficient to explain the diversity of downstream effects conferred by GPCR transit through CCPs.

Here we show that release of the β-arrestin CT is only one mode by which GPCR binding can trigger the endocytic activity of β-arrestins. We resolve a second mode that is clearly distinct from the canonical CT-dependent mode because it does not require the β-arrestin CT whatsoever. We further show that GPCRs exhibit selectivity in triggering one or the other endocytic mode, and that each mode is differentially coupled to the GPCR / β-arrestin interface to produce opposing effects on the net signaling output of receptors.

## Results

### The β-arrestin C-terminus is dispensable for β2AR internalization

The ability of the β-arrestin CT to associate with CCP components and influence GPCR endocytosis has been clearly established. However, to our knowledge, whether the β-arrestin CT is necessary for GPCR endocytosis has not been explicitly tested. To assess this, we pursued a genetic rescue strategy using CRISPR-engineered cells lacking both β-arrestins (βarr1/2 DKO) (O’Hayre et al. 2017) and asked if β-arrestin-2 (βarr2) is sufficient to rescue regulated endocytosis of the β2-adrenergic receptor (β2AR), a prototypic β-arrestin-dependent cargo. We verified that our experiments were carried out in an expression regime in which receptor internalization remains phosphorylation dependent (Extended Data Fig. 1g), and began by examining agonist-induced clustering of receptors into CCPs on the plasma membrane as an essential intermediate step in the endocytic mechanism. We verified by total internal reflection fluorescence (TIRF) microscopy that β2ARs localize diffusely on the plasma membrane in βarr1/2 DKO cells, regardless of whether cells were exposed to the adrenergic agonist isoproterenol (Iso, Fig. 1a). However, after expression of recombinant βarr2 (βarr2-EGFP), Iso induced β2ARs to rapidly transition from a diffuse distribution (Fig. 1a) to co-clustering together with βarr2 in CCPs (Fig. 1b).

**Fig. 1.**
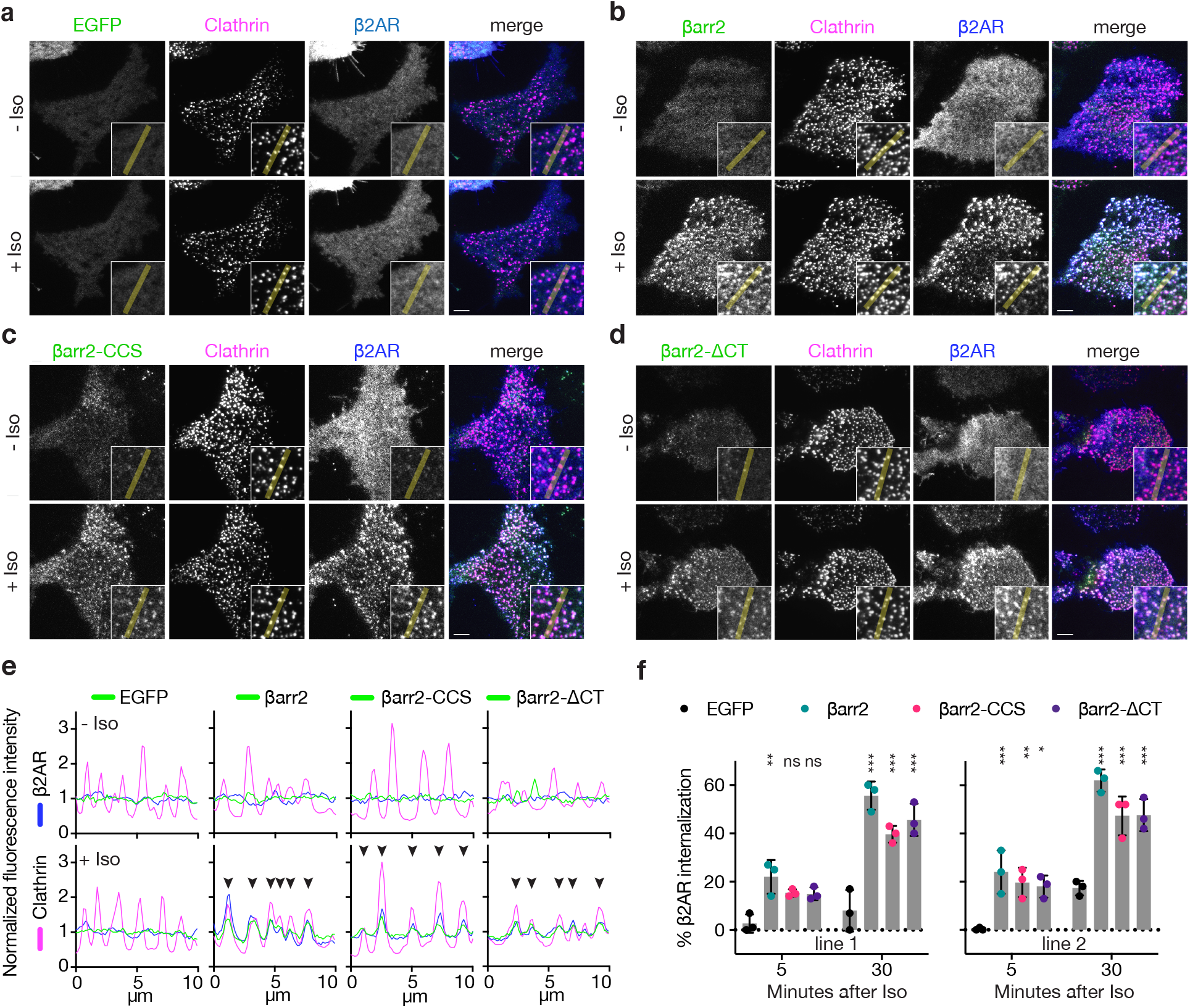
Known endocytic motifs in βarr2 are dispensable for β2AR clustering and endocytosis. **a-d**, Representative live-cell TIRF microscopy images of βarr1/2 double knockout HEK293 cells co-expressing clathrin light chain-DsRed (magenta) and FLAG-tagged β2AR (blue) with either EGFP (a), βarr2-EGFP (b), β arr2-CCS-EGFP (c), or βarr2-ΔCT-EGFP (d) (all in green) and pre- and post-stimulation with 10 μM isoproterenol (Iso). Scale bars at 5 μm. **e**, Representative fluorescence intensity profiles from line scans shown in insets from a-d. **f**, Internalization of FLAG-tagged β2AR co-expressed with either EGFP (black), wild type βarr2-EGFP (green), βarr2-CCS-EGFP (pink), or βarr2-ΔCT-EGFP in two clonal lines of βarr1/2 DKO HEK293s at 5- and 30-minutes post-stimulation with 10 μM isoproterenol (Iso). Data shown as mean ± s.d. for n = 3 independent experiments.. Significance was determined by two-way ANOVA (df = 3, F = 24.48) with Tukey’s multiple comparisons test against the negative control (EGFP) for each time point (ns P ≥ 0.05, * P < 0.05, ** P < 0.01, *** P < 0.001). All data shown is from three independent experiments.

Surprisingly, genetic rescue of βarr2-driven clustering was observed even after clathrin and AP2 binding elements in the β-arrestin CT were disrupted using previously defined point mutations (βarr2-CCS construct) (Kelsie Eichel et al. 2018). Confirming this, βarr2 rescued clustering and internalization after both elements were fully removed by truncation (βarr2-ΔCT construct) (Fig. 1c-e). The latter result was verified in two independent βarr1/2 DKO cell lines (Fig. 1f) and it was not limited to the β2AR (Extended Data Fig. 1b-f). Moreover, CT-independent trafficking was not unique to β-arrestin-2, as a β-arrestin-1 mutant lacking the CT (βarr1-ΔCT) also promoted Iso-induced β2AR internalization (Extended Data Fig. 1h). Taken together, these observations indicate that the β-arrestin CT is indeed not required for β-arrestin to drive clustering of GPCR/β-arrestin complexes on the cell surface or promote subsequent endocytosis of complexes. As both clustering and endocytosis mediated by βarr2-ΔCT remained agonist-dependent, these results suggest that β-arrestins contain additional endocytic determinant(s) capable of mediating GPCR-triggered endocytic activity in the absence of the β-arrestin CT.

### A discrete endocytic determinant in the β-arrestin C-lobe

To investigate the nature of this additional endocytic activity, we took advantage of the fact that visual arrestin (v-arr) naturally fails to support GPCR endocytosis (Moaven et al. 2013) and asked if we could generate a v-arr / βarr2 chimera that contains the C-terminus of v-arr but retains the ability to promote GPCR endocytosis due to the addition of other βarr2-derived sequence(s) (Extended Data Fig. 2a,b). We tested this using the same genetic rescue strategy. To focus specifically on effects downstream of GPCR binding, we measured internalization using a chimeric mutant receptor (β2V2R chimera) that has higher affinity for binding arrestins than the β2AR (Oakley et al. 2000, 2001). We were indeed able to generate such a chimeric mutant arrestin construct, here called ChiA, which rescued agonist-induced internalization of receptors nearly as strongly as wild type βarr2 despite its entire C-terminus being derived from v-arr (Fig. 2a).

**Fig. 2.**
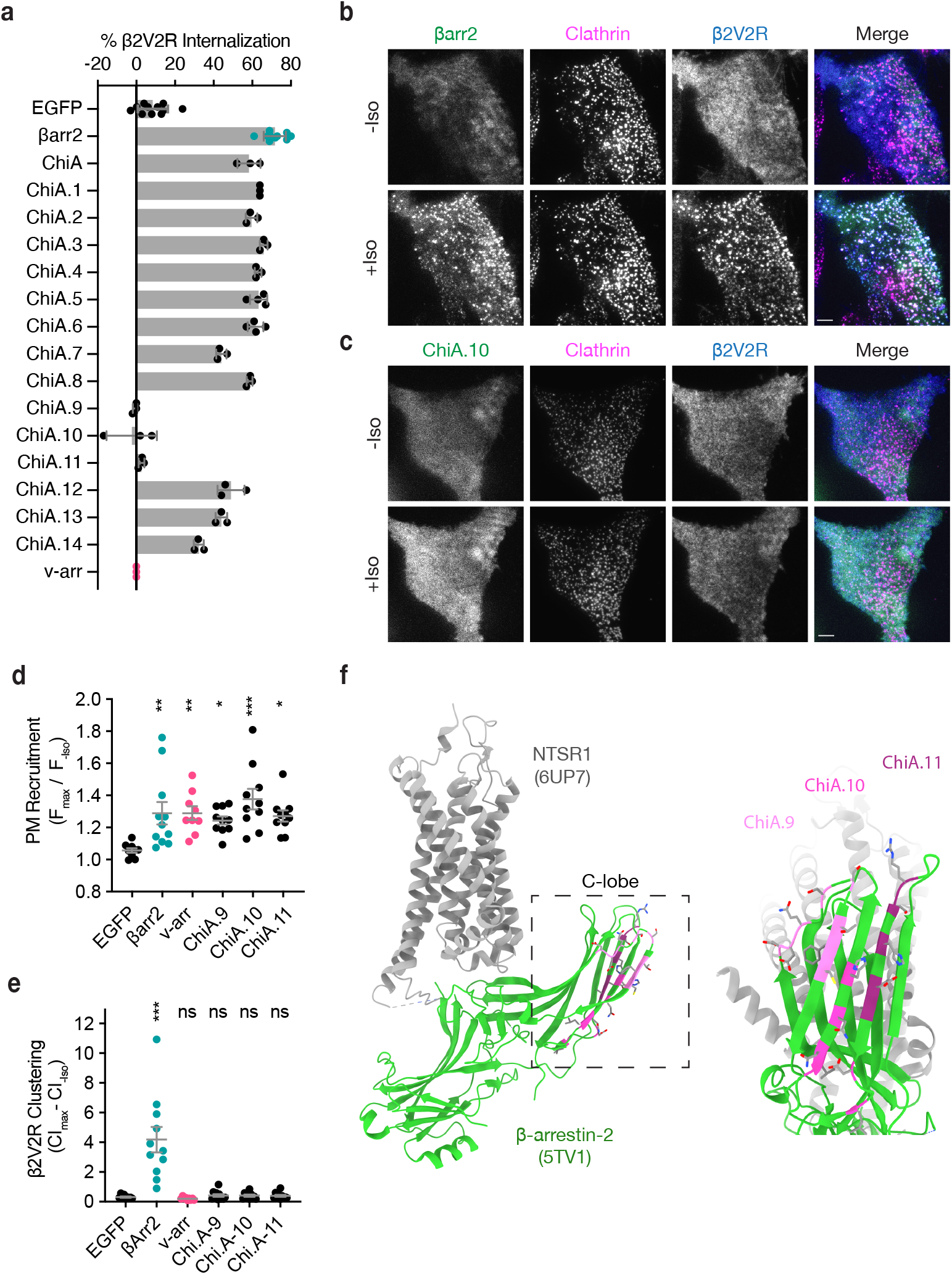
Identification of the βarr2 C-lobe base (CLB) **a**, Internalization of β2V2R after 30 minutes of 10 μM isoproterenol stimulation in βarr1/2 DKO HEK293s co-expressing the indicated construct (n ≥ 3 independent experiments, line is mean, error bars are s.d., each dot is an average of three technical replicates). **b, c**, Representative TIRF microscopy images of cells expressing β2AR (blue) and clathrin-light-chain-dsRed (magenta) with either wild-type βarr2-EGFP (b) or an example of one of the three internalization defective chimeras, ChiA.10-EGFP (c), pre- and post-stimulation with 10 μM isoproterenol (Iso). Scale bars are 5 μm. **d**, Plasma membrane recruitment of the indicated EGFP-tagged proteins (see methods) in response to stimulation with 10 μM isoproterenol. **e**, Maximum clustering index (CI, see methods) of plasma membrane β2V2R after treatment with 10 μM Iso. For **d** and **e**, each dot represents an individual cell. Data are shown as mean ± s.e.m. (n ≥9 cells). Significance was determined by ordinary one-way ANOVA (df = 5 for both, F = 22.21 and 4.531, respectively) with Dunnett’s multiple comparison test against negative control (EGFP) (ns P ≥ 0.05, * P < 0.05, ** P < 0.01, *** P < 0.001). **f**, Location of mutations unique to ChiA.9-11 (shades of pink and purple) in an active state structure of β-arrestin-2 (5TV1, green) (Chen et al. 2017) fit to the NTSR1/βarr1 structure (6UP7, gray) (Huang et al. 2020) (βarr1 not shown) and the same model rotated and zoomed to the cytoplasmic face of the C-lobe. All data shown is from at least three independent experiments.

To identify sequence(s) responsible for the endocytic activity of this chimeric arrestin protein, we systematically replaced small sections of βarr2-derived sequence in ChiA to the corresponding sequence in v-arr (ChiA.1-14). Of 14 chimeras tested, we found three– ChiA.9, 10, and 11– that failed to rescue internalization (Fig. 2a). TIRF imaging indicated that all these internalization-defective chimeras were recruited to the plasma membrane in response to Iso addition, and to a similar degree as wild type βarr2.

However, they localized diffusely and failed to cluster receptors in CCPs. This was evident visually (Fig. 2b, c) and quantified by fluorescence intensity measurement (Fig. 2d, e). These data suggest that each of the mutations specifically interferes with the clustering and endocytic function of β-arrestin-2 by preventing accumulation in CCPs, but without affecting receptor-triggered recruitment to the plasma membrane. The endocytosis-blocking mutations mapped to a contiguous region of β-arrestin-2, located at the cytoplasmic face of the C-lobe and opposite the receptor binding interface, which we called the β-arrestin C-lobe base (CLB, Fig. 2f).

### The C-lobe determinant is essential for β2AR internalization

We next investigated the contributions of mutating the CLB or CT on the natural endocytic activity of βarr2. We focused on the central β-strand in the CLB of βarr2 and introduced mutations corresponding to residues present in v-arr (D205S, L208I, L215I, N216P, N218T, and H220A), while leaving the βarr2 CT unchanged. The resulting mutant construct, βarr2-CLB, was strongly recruited to the plasma membrane after Iso-induced activation and clustered together with β2AR in CCPs (Fig. 3b). When the CLB was mutated in combination with the CT (βarr2-CLB,ΔCT mutant), agonist-induced recruitment to the plasma membrane was retained but βarr2-CLB,ΔCT was recruited diffusely and failed to promote surface clustering of β2ARs (Fig. 3d). More detailed inspection of TIRF microscopy images revealed that mutating either the CLB or CT produced a partial reduction in clustering relative to wild type βarr2, whereas clustering was abolished in the double mutant (Fig. 3e). We verified this specific effect on clustering using a previously established quantitative metric (Fig. 3f,g) (Kelsie Eichel et al. 2018), despite all of the constructs being similarly recruited to the plasma membrane (Fig. 3h). Together, these results indicate that the β-arrestin CT and CLB promote β2AR clustering in an additive manner. As βarr2-CLB,ΔCT failed to cluster at all, the results also indicate that the CT and CLB, together, fully account for the GPCR-triggered clustering activity of βarr2.

**Fig. 3.**
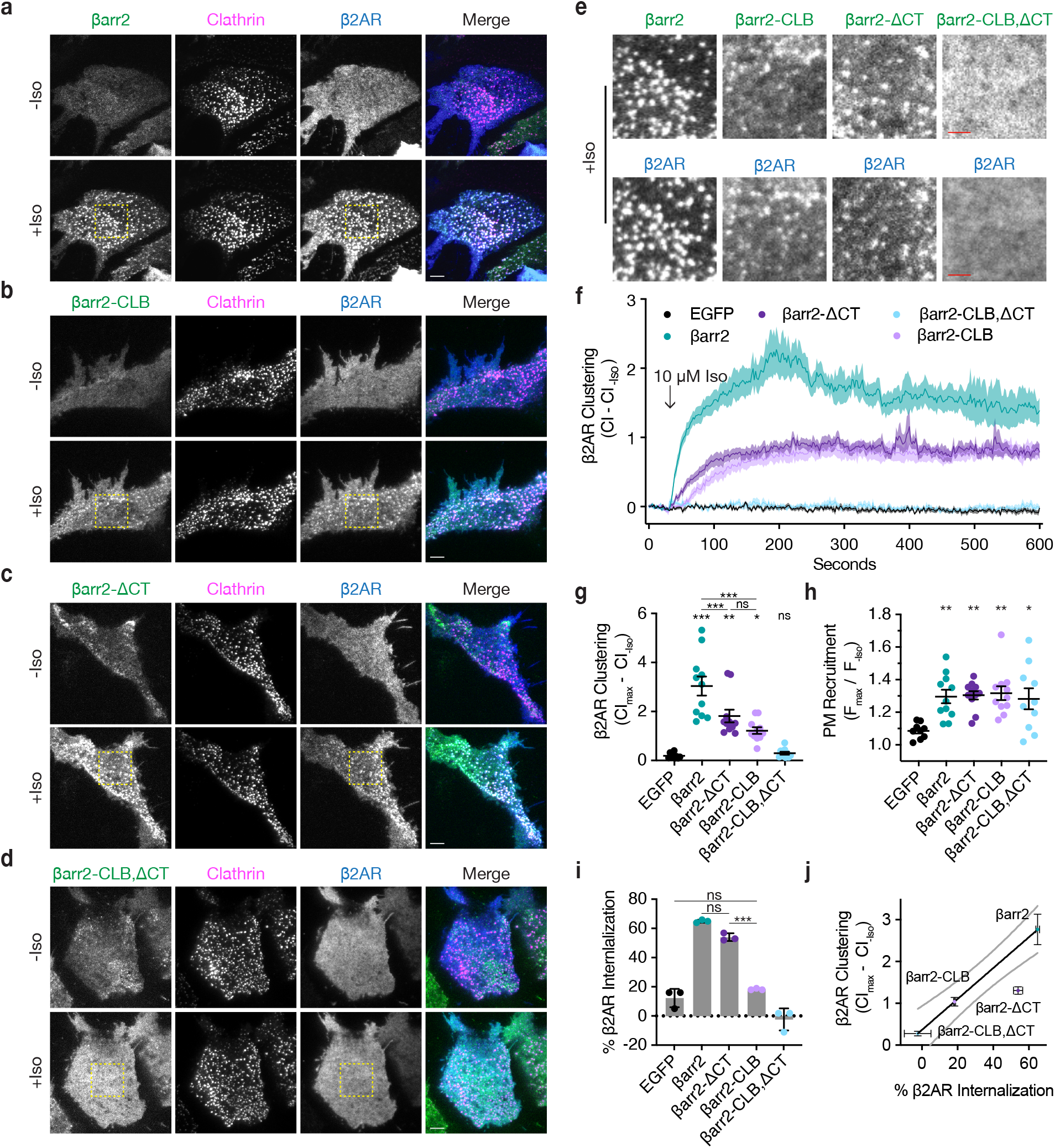
βarr2 CT is not sufficient for β2AR internalization. **a**-**d**, Representative live-cell TIRF microscopy images of βarr1/2 double knockout HEK293s co-expressing clathrin light chain-DsRed (magenta) and FLAG-tagged β2AR (blue) with either EGFP-tagged βarr2 (a), β arr2-CLB (b), βarr2-ΔCT (c), or βarr2-CLB,ΔCT (d) (all in green) pre- and post- stimulation with 10 μM isoproterenol (Iso). Scale bars represent 5 μm. EGFP condition not shown (see fig. 1A for example). **e**, Zoomed images corresponding to boxes in panels a-d for βarr2 and β2AR images. Scale bars (red) represent 2.5 μm. **f**, β2AR clustering index (CI, see methods) pre- and post-stimulation with 10 μM Iso over ten minutes. **g**, Max plasma membrane recruitment of the indicated EGFP-tagged proteins in response to treatment with 10 μM Iso. **h**, Max clustering index (CI) of β2AR calculated from within the first 300 seconds of (f) and normalized to clustering index prior to Iso treatment. For **f, g**, and **h**, data shown as mean ± s.e.m. (n ≥ 9 cells, represented as dots in (g) and (h)) **i**, Internalization of β2AR when co-expressed with the indicated EGFP-tagged proteins (n = 3, each dot is an average of three technical replicates) in βarr1/2 DKO HEK293 cells. **j**, Correlation between β2AR clustering and internalization. Solid line is a simple linear regression fit to βarr2 and βarr2-CLB,ΔCT. (R2 = 0.69, dashed lines = 95% CI, vertical error = s.e.m. and horizontal error = std. dev.). For **g, h, i**, significance was determined by ordinary one-way ANOVA (df = 4 for all, F = 21.32, 4.828, and 117.6, respectively) with Tukey’s test for multiple comparisons (ns P ≥ 0.05, * P < 0.05, ** P < 0.01, *** P < 0.001). All data shown is from at least three independent experiments.

Despite the ability of the CT and CLB to promote β2AR clustering to a similar degree (Fig. 3g), their effects on the subsequent internalization of receptors differed considerably. Mutating the CT only slightly reduced the ability of βarr2 to drive internalization of β2ARs, whereas mutating the CLB caused a pronounced reduction– suppressing the measured internalization of receptors to a level similar to that of the EGFP negative control (Fig. 3i). The same effect was observed when the CLB was similarly mutated in βarr1 (βarr1-CLB, Extended Data Fig. 3a), indicating that this determinant functions similarly in both β-arrestins. Further, the activity of the CLB was impaired by introducing a single point mutation, N218T, into otherwise wild type βarr2, changing a single solvent-exposed residue in the CLB to the corresponding residue in v-arr (Extended Data Fig. 3b,c). Taken together (Fig. 3j), these data suggest that both the CLB and CT contribute to the total endocytic activity of β-arrestin, and each promotes agonist-dependent clustering of receptors on the cell surface. However, the CLB, rather than CT, is the primary determinant in β-arrestin required for the efficient internalization of β2ARs.

### GPCRs differ in the degree to which they utilize each endocytic determinant

To assess the role of each determinant more broadly, we examined the effect of individually mutating the CT (βarr2-ΔCT) or CLB (βarr2-CLB) on agonist-induced internalization across a number of β-arrestin-dependent GPCRs. In an effort to include receptors differing in engagement at the GPCR / β-arrestin interface, we included three naturally occurring ‘class A’ GPCRs (β2AR, μ-opioid receptor or μOR, and κ-opioid receptor or κOR) that associate with β-arrestins relatively weakly and three ‘class B’ GPCRs (V2 vasopressin receptor or V2R, M2 muscarinic receptor or M2R, and neurotensin receptor type 1 or NTSR1) that bind β-arrestins more strongly (Oakley et al. 2000, 2001). In addition, we included a chimeric GPCR containing the β2AR-derived transmembrane core and V2R-derived cytoplasmic tail (β2V2R) that is widely used as an experimental model of a class B GPCR.

Removing the β-arrestin-2 CT had little effect on agonist-induced internalization of the class A GPCRs, but significantly inhibited internalization of the naturally occurring class B GPCRs. By contrast, mutating the CLB inhibited internalization of all receptors tested, although the magnitude of this effect varied depending on the receptor (Fig. 4a). Together, these results indicate that both the CT and CLB are broadly utilized endocytic determinants, but that individual GPCRs vary significantly in the degree to which their endocytosis depends on each.

**Fig. 4.**
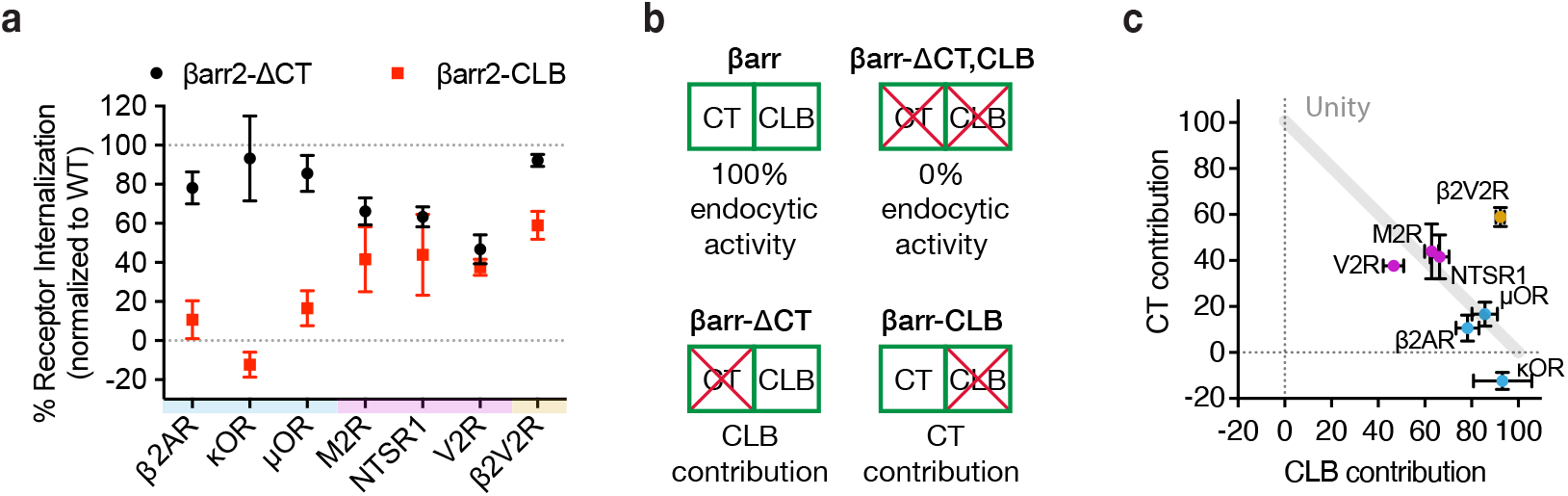
GPCRs selectivity utilize the βarr2 CLB and CT for endocytosis. **a**, Internalization of the CT (black) and CLB (red) mutants normalized to wild type βarr2 for each receptor shown 30 minutes after agonist addition (see Extended Data Fig. 4). Each dot is the mean of three independent experiments ± standard deviation. Shading indicates whether receptors are naturally occuring class a (blue), class b (magenta), or engineered class b (gold). **b**, Schematic summarizing the conceptual basis for estimating contributions of the CT and CLB. Contribution of each determinant within βarr2 is defined by subtracting internalization measured in the negative control (EGFP) from βarr2, βarr2-ΔCT, βarr2-CLB and dividing the resulting values by control subtracted wild-type (βarr2) value. **c**, Contribution to total endocytic activity of each determinant plotted as x and y coordinates for each receptor from panel (a). Unity is defined as 100% endocytic activity when individual activities are summed. Dot color corresponds to the typology described for panel (a). All experiments were performed in βarr1/2 DKO HEK293 cells.

Despite GPCR-specific differences in reliance on the CT relative to CLB, mutating both the CT and CLB in combination (βarr2-CLB,ΔCT) abolished the endocytic activity of β-arrestin-2 for all of the GPCRs (Extended Data Fig. 4a-f). This indicates that the CT and CLB are sufficient to fully account for the GPCR-triggered endocytic activity of β-arrestin-2. Accordingly, we used the effect of mutating each determinant individually to estimate the relative contribution of the other (non-mutated) determinant (Fig. 4b).

A roughly linear relationship between the relative contribution of each determinant was observed for all of the naturally occurring GPCRs included in our panel, with the contribution of both determinants summing to approximately 100% (Fig. 4c). When compared in this way, receptors clustered according to differences in overall stability, as defined by the ‘class A/B’ classification scheme. The class A GPCRs (β2AR, κOR, μOR), defined by binding β-arrestins relatively weakly or transiently, were found to primarily utilize the CLB with little utilization of the CT. The class B GPCRs (V2R, M2R, NTSR1), defined by binding β-arrestins more strongly or stably, utilized both determinants in an additive manner.

The β2V2R chimera is considered a class B GPCR due to the V2R-derived cytoplasmic tail conferring enhanced binding to β-arrestin. This chimeric receptor also utilized both the CLB and CT for endocytosis, providing further support for the concept that the overall strength of GPCR/β-arrestin binding impacts the utilization of endocytic determinants. Unlike the other receptors, however, the β2V2R departed from the linear relationship because the estimated contribution of each determinant summed to more than 100%. This suggests that this engineered GPCR can strongly utilize either the CT or CLB for endocytosis in a semi-redundant manner.

We further noted that, when compared to the β2AR, the β2V2R departed from the unity line primarily due to a stronger ability to use the β-arrestin CT for endocytosis (Fig. 4c). This is interesting because the β2V2R differs from the wild type β2AR mainly in its tail interaction with β-arrestin mediated by V2R-derived tail sequence. Relative to the V2R, the β2V2R departed from the linear relationship primarily due to stronger utilization of the β-arrestin CLB. This is interesting as β2V2R is thought to differ from the wild type V2R primarily in its core interaction with β-arrestin. Accordingly, we speculate that GPCRs differ in their ability to utilize the CT or CLB for endocytosis according to differences in the interactions that they make with β-arrestin through the receptor tail and core, respectively.

### Each endocytic determinant is oppositely coupled to the net signaling output of endogenous β2ARs

Having observed that differences in the GPCR/β-arrestin complex affect utilization of the CT relative to CLB, we next asked if each discrete endocytic determinant conversely influences the GPCR/β-arrestin interaction. To test this, we returned to the β2AR as a prototype and investigated how mutating the CT or CLB affects β2AR/βarr2 complex formation in living cells.

We first assessed complex formation using a real-time protein complementation assay based on an engineered split luciferase enzyme (NanoBiT) (Dixon et al. 2016; Inoue et al. 2019). Iso produced a concentration-dependent increase in the interaction between the β2AR and wild type β-arrestin-2. Similar results were obtained after mutating the CT and CLB individually or in combination, verifying that neither determinant is required for receptor/β-arrestin complex formation. However, we noted a trend suggesting additional specificity: Mutating the β-arrestin-2 CLB (βarr2-CLB) appeared to decrease the agonist potency (reported as logEC50 ± 95%CI) for recruitment (−7.3 ± 0.1), whereas mutating the CT (βarr2-ΔCT) increased it when compared to wild-type (−6.8 ± 0.2 and −7.1 ± 0.1, respectively), and mutating both determinants in combination (βarr2-CLB,ΔCT) trended toward an intermediate effect (−7.1 ± 0.3) (Fig. 5a). We also noted that single mutation of the CLB slowed β2AR/βarr2 association relative to that observed for wild type protein (Tau95%CI = 81 - 108 seconds and 23 - 56, respectively), whereas the double (CLB and CT) mutation produced kinetics indistinguishable from wild type (Tau95%CI = 31 - 37 seconds, Fig. 5b). These results indicate that neither the CT nor CLB is required for β-arrestin to associate with receptors, but suggest that each determinant influences the interaction in different ways.

**Fig. 5.**
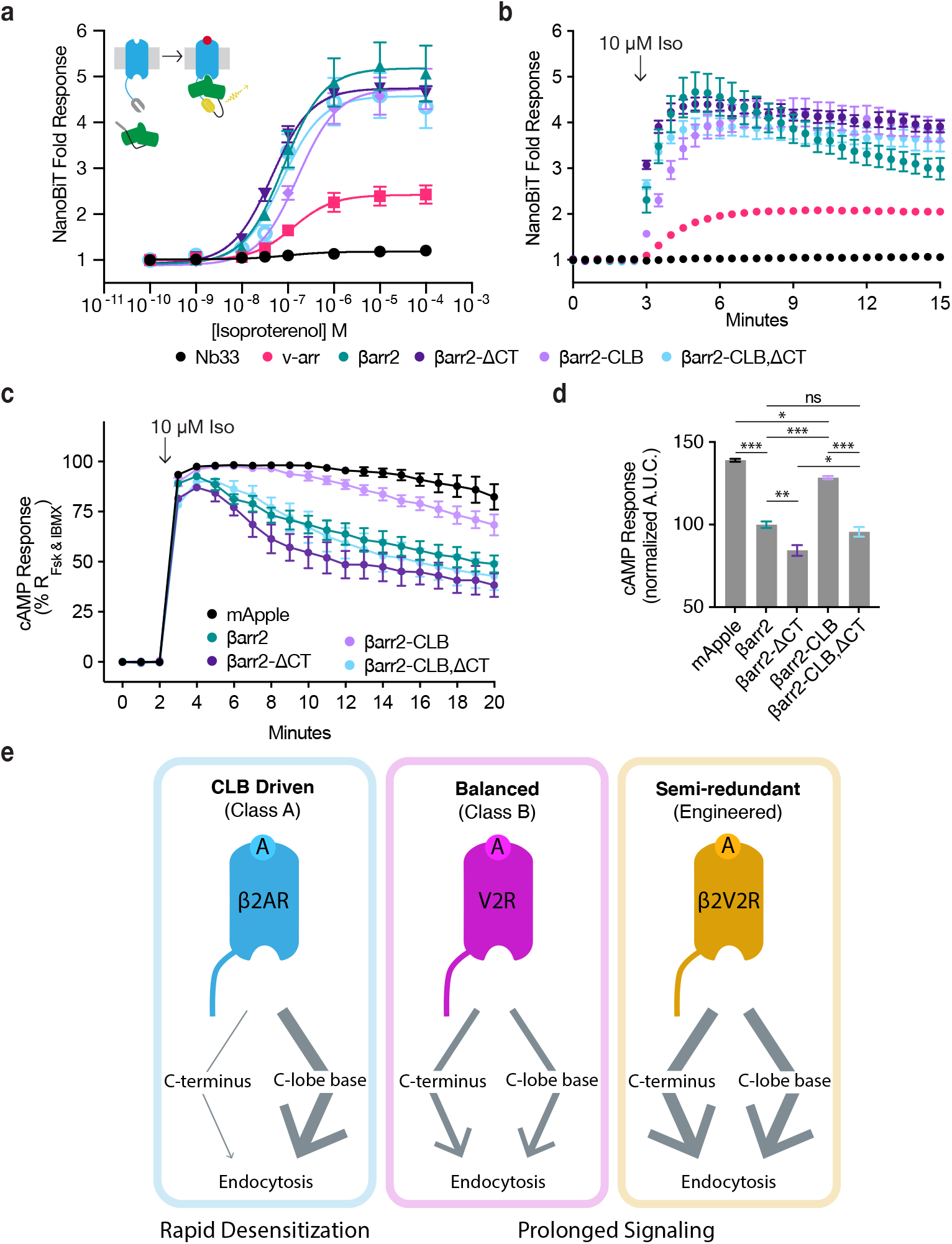
CLB and CT determinants reveal two allosteric paths from GPCRs to the endocytic network. **a, b**, Direct NanoBiT luciferase complementation of β2AR-LgBiT and SmBiT-tagged: Nb33 (a μOR receptor specific nanobody, black), visual arrestin (pink), wild type β-arrestin-2 (green), CT mutant (dark purple), CLB mutant (light purple), or double mutant (cyan) measured as an end point across a range of isoproterenol (Iso) concentrations (a) and kinetically (b) pre- and post-stimulation with 10 μM Iso. Dose response curves were generated with three-parameter nonlinear fit (R2 = 0.94-0.99). **c**, Endogenous β2AR cAMP response after stimulation with 10 μM Iso measured by a genetically encoded fluorescent cAMP biosensor, cADDis, and normalized to the response elicited by simultaneous treatment with 10 μM forskolin (Fsk) and 300 μM 3-isobutyl-1-methylxanthine (IBMX). **d**, Area under the curve calculated from panel (c). All data shown as mean ± s.e.m. from three independent experiments performed in βarr1/2 DKO HEK293 cells. Significance was determined by an ordinary one-way ANOVA (df = 4, F = 112.1) with Tukey’s multiple comparisons test. ns P ≥ 0.05, * P < 0.05, *** P < 0.001. **e**, Diagram of proposed model involving two differentially utilized allosteric paths from GPCRs through β-arrestins to promote endocytosis. Class A GPCRs (blue), exemplified by the β2AR, primarily utilize the CLB to drive endocytosis while Class B GPCRs, exemplified by V2R (magenta) and β2V2R (gold) utilize both determinants. Arrows represent the proposed allosteric paths linking the GPCR / β-arrestin interface to the β-arrestin / CCP interface, explaining how the CLB-dependent endocytic mode is coupled to rapid desensitization of receptor signaling while the CT-dependent mode enables prolonged signaling.

More pronounced differences were observed when we assessed β2AR/β-arrestin complex formation functionally by assessing desensitization of the endogenous β2AR-elicited cAMP response. We verified a lack of β-arrestin-mediated desensitization in βarr1/2 DKO cells, as indicated by persistently elevated cAMP in the continued presence of agonist, and rescue of desensitization by recombinant wild type βarr2 (Fig. 5c, d). We also observed rescue of desensitization by the βarr2-CLB,ΔCT double mutant construct, despite this construct having no detectable endocytic activity, as well as rescue after single mutation of the β-arrestin CT. Remarkably, individually mutating the CLB abrogated the ability of βarr2 to rescue the desensitization response (Fig. 5c, d). Moreover, this signaling effect was phenocopied by making the same single-residue exchange (N218T) in the otherwise wild type CLB (Extended Data Fig. 5a, b). Together, these results suggest that the discrete endocytic determinants in the β-arrestin CLB and CT, while not known to directly contact the receptor and not essential for GPCR/β-arrestin complex formation, are each coupled to the receptor/β-arrestin interface and produce opposing effects on the net signaling output of receptors.

## Discussion

Prior to this work, the ability of GPCRs to trigger the endocytic adaptor activity of β-arrestin was assumed to rely entirely on release of the β-arrestin CT. Revising this view, we show that the β-arrestin CT, while clearly capable of promoting GPCR endocytosis, is not necessary for this process. We define the CLB as a discrete determinant in β-arrestin that is critical for the endocytic activity of β-arrestin, and which can operate in the absence of the CT. We further show that the relative contribution of the β-arrestin CT and CLB in mediating the endocytic activity of β-arrestin varies depending on the GPCR bound, with class A receptors primarily dependent on the CLB and class B GPCRs dependent on the CT and CLB to a similar degree. By focusing on the β2-adrenergic receptor as a prototypic class A GPCR, we additionally show that the β-arrestin CT can promote receptor clustering into CCPs despite having little ability to drive the subsequent endocytosis of receptors. Moreover, we show that mutations which selectively disrupt each of the endocytic determinants produce opposing effects on the net signaling output of receptors. In sum, these results support a model in which the endocytic activity of β-arrestins is triggered flexibly by GPCRs through distinct (CT and CLB-dependent) biochemical modes that differentially attenuate or prolong receptor signaling (Fig. 5e).

Only two β-arrestins regulate hundreds of GPCRs and it is increasingly clear that β-arrestins function as highly flexible regulators of GPCR signaling, with the ability to produce a variety of downstream functional effects depending on receptor-specific differences communicated across the GPCR / β-arrestin interface. The present results extend this concept to flexibility and diversity at the β-arrestin / CCP interface. Our finding that mutations which disrupt either the CT or CLB produce different effects on endogenous β2AR signaling suggest that each endocytic mode, in addition to being triggered in a GPCR-specific manner, is coupled to opposing effects on the net signaling output of receptors.

Beyond providing new insight into the cell biology of GPCRs and β-arrestins, we believe that our results have implications for understanding endocytic adaptor functions more generally. The best studied among these is AP2, a protein complex that is essential for CCP nucleation and co-assembles with clathrin. AP2 switches from a ‘closed’ to ‘open’ state triggered by the binding of PIP2 and endocytic cargo (Kovtun et al. 2020). β-arrestins are structurally distinct from AP2 and also differ functionally, as they are not required for CCP formation and associate with CCPs primarily after assembly (Puthenveedu and von Zastrow 2006; Santini, Gaidarov, and Keen 2002). Nevertheless, β-arrestins also switch from ‘closed’ to ‘open’ states triggered by binding GPCR cargo and aided, in many cases, by the binding of PIP2 (Kang, Tian, and Benovic 2014; Shukla et al. 2013; Huang et al. 2020; Gaidarov et al. 1999). The present results provide evidence that β-arrestin can switch into more than one active or ‘open’ state, and do so selectively depending on the GPCR bound. To our knowledge, these findings provide the first example of an endocytic adaptor protein capable of undergoing cargo-specific mode switching. The existence of such a flexible relationship between cargo binding and functional activation of the adaptor protein suggests that adaptor proteins have the capacity to communicate more nuanced information to the CCP than the simple presence or absence of a cognate cargo. In addition, these findings raise multiple new questions about the biochemical and structural basis for the discrete active or ‘open’ states of β-arrestin that we resolve here according to functional differences.

### Ideas and Speculation

While the present results provide strong evidence for the existence of discrete endocytic modes differing in dependence on the β-arrestin CT and CLB, they leave unresolved the biochemical details of the latter mode. In contrast to the β-arrestin CT, for which relevant CCP-associated interacting partners are well known, how the β-arrestin CLB engages CCPs remains to be determined. We speculate that the discrete function of the CLB depends on engagement with partners other than the clathrin terminal domain and the β-appendage of AP2, which are key domains interacting with the β-arrestin CT. The relatively understudied C-lobe engages multiple additional proteins and we note that two of these, PIP5K1A (Nelson et al. 2008; Jung et al. 2021) and PDE4D5 (Baillie et al. 2007; Perry et al. 2002), have binding sites which overlap the CLB and are already known to affect endocytosis and GPCR desensitization. However, neither protein is currently known to be enriched in CCPs, which is presumably required for the CLB to drive GPCR-triggered accumulation in CCPs without the CT as we show. Moreover, the CCP lattice associates with more than 50 additional proteins, some only transiently during CCP maturation, and a number with known regulatory effects (Mettlen et al. 2018; Merrifield and Kaksonen 2014; Kirchhausen, Owen, and Harrison 2014; Traub 2011). Thus, significant further study will be required to determine whether and how these proteins, and/or other candidates yet to be identified, mediate the signaling and trafficking functions of the CLB as identified here.

### Materials and Methods

### Cell culture, expression constructs, and transfections

βarr1/2 double knockout HEK 293A cell lines, generously provided by Asuka Inouye and Silvio Gutkind (O’Hayre et al. 2017), were cultured in complete growth Dulbecco’s modified Eagle’s medium (DMEM, Life Technologies, 11965118) supplemented with 10% fetal bovine serum (UCSF Cell Culture Facility). Cell cultures were verified to be free of mycoplasma contamination by enzymatic assay (MycoAlert, Lonza, LT07-318). Transfections were carried out using Lipofectamine 2000 (Thermo, 11668019) according to the manufacturer’s protocol. Cells were transfected 24-48 h before experiments.

All receptor constructs were N-terminally FLAG-tagged. The human β1AR, β2AR, V2R, μOR, and κOR were previously described (Cao et al. 1999; Temkin et al. 2011; Chu et al. 1997). NTSR1 was a generous gift from Brian Kobilka (Huang et al. 2020). The β2AR–V2R chimera was a generous gift from Marc Caron (Oakley et al. 1999). M2R was previously generated in the lab by James Hislop. The β2AR-3S was previously described (Hausdorff et al. 1991).

β-arrestin-2–GFP and β-arrestin-2–mApple were previously described (Barak et al. 1997; K. Eichel, Jullié, and von Zastrow 2016). β-arrestin-1–EGFP was generated by PCR amplifying from β-arrestin-1–mVenus, which was a gift from R. Sunahara (University of California, San Diego), and subcloned into EGFP-N1.

Visual arrestin–EGFP was generated by synthesis of bovine visual arrestin (Twist Biosciences) and directly subcloned into EGFP-N1 using InFusion (Takara). β-arrestin-2-CCS–EGFP was previously described (Kelsie Eichel et al. 2018). β-arrestin-2-ΔCT–EGFP was generated by PCR amplifying from β-arrestin-2– EGFP and subcloning into EGFP-N1 using InFusion HD (Takara Bio). β-arrestin-2-SmBiT constructs were generated by removing EGFP from βarr2, βarr2-ΔCT, βarr2-CLB, βarr2-ΔCT,CLB constructs by digestion with ApaI and XbaI followed by insertion of SmBiT by two rounds of PCR followed by InFusion HD (Takara Bio).

Visual arrestin and β-arrestin-2 EGFP-tagged chimeras were generated by synthesis of the template chimera, ChiA, (Twist Bioscience) and subcloned into EGFP-N1. Subsequent chimeras, ChiA.1-14, were generated by PCR and InFusion HD (Takara Bio).

Clathrin–dsRed was previously described (Merrifield et al. 2002).

Nb33-SmBiT was generated in the lab by Joy Li.

### Live cell TIRF microscopy imaging

TIRF microscopy was performed at 37 °C using a Nikon Ti-E inverted microscope equipped for through-the-objective TIRF microscopy and outfitted with a temperature-, humidity- and CO2-controlled chamber (Okolab). Images were obtained with an Apo TIRF 100 X, 1.49 numerical aperture objective (Nikon) with solid-state 405, 488, 561 and 647 nm lasers (Keysight Technologies). An Andor iXon DU897 EMCCD camera controlled by NIS-Elements 4.1 software was used to acquire image sequences every 2 s for 10 min. βarr1/2 double knockout HEK293s were transfected as indicated according to the manufacturer’s protocol 48 h before imaging and then plated on poly-L-lysine (0.0001%, Sigma) coated 35-mm glass-bottom culture dishes (MatTek Corporation) 24 h before imaging. Cells were labeled with monoclonal FLAG antibody (M1) (1:1000, Sigma F-3040) conjugated to Alexa Fluor 647 dye (Life Technologies) for 10 min at 37 °C before imaging, washed, and imaged live in DMEM without phenol red (UCSF Cell Culture Facility) supplemented with 30 mM HEPES, pH 7.4 (UCSF Cell Culture Facility). Cells were treated by bath application of isoproterenol at the indicated time. At least three independent experiments were performed for all live-cell microscopy.

### TIRF microscopy image analysis

Quantitative image analysis was performed on unprocessed images using ImageJ and Fiji software (Schindelin et al. 2012; Schneider, Rasband, and Eliceiri 2012). To quantify change in β-arrestin fluorescence over time in TIRF microscopy images, which was reported as plasma membrane recruitment, fluorescence values were measured over the entire time series in a region of interest (ROI) corresponding to the cell. Fluorescence values of the ROI were normalized to fluorescence values before agonist addition. Minimal bleed-through and photobleaching was verified using single-labeled and untreated samples, respectively. Line scan analysis of receptor, β-arrestin, or clathrin fluorescence was carried out using the Fiji plot profile function to measure pixel values from the five pixel wide lines shown. Clustering index was determined using the skew statistical measurement applied to fluorescence intensity values of M1-Alexa647 labeled receptor pixels in a ROI corresponding to the cell as has been previously described (Kelsie Eichel et al. 2018).

### Flow cytometry internalization assays

All internalization assays were performed with βarr1/2 DKO HEK293 cells that were transfected in 6 cm dishes according to the manufacturer’s protocol 24 hours before beginning the assay with 1000 ng of GPCR DNA paired with 400-800 ng of βarr (or one of its mutants) DNA. At least three hours before drug treatment cells were lifted using TrypLE Express (Thermo Scientific), a dissociation reagent that leaves extracellular epitopes intact, resuspended in complete media, transferred to 12-well plates in triplicate, and incubated under standard culture conditions. Cells were then treated with agonist(s) (see figure legends) for the indicated period and placed on ice to stop trafficking. Cells were washed once with ice cold PBS supplemented with 2 mM CaCl2 followed by labeling of FLAG-tagged surface receptors with M1 antibody (Sigma Aldrich, F3040) conjugated to Alexa Fluor 647 (Thermo Scientific) for 30 minutes at 4°C while gently shaking. Surface staining of receptors was measured using a CytoFlex (Beckman Coulter) with gates set for single cells expressing EGFP. Examination of unstimulated cells verified that fluorescence from either EGFP or surface receptors was similar across conditions within each GPCR tested. Percent internalization was calculated by taking the mean M1-647 fluorescence for the agonist treated cells and dividing it by the same measure for the corresponding unstimulated cells, subtracted from one, and multiplied by 100. At least three independent experiments were performed for all internalization assays.

### NanoBiT complementation

β-arrestin-1/2 double knockout HEK293s were plated in 6-well dishes, transfected with β2AR-LgBiT (200 ng per well of a six-well plate) paired with one of the following: Nb33-EGFP-SmBiT, arr1-SmBiT, βarr2-SmBiT, βarr2-ΔCT-SmBiT, βarr2-CLB-SmBiT, or βarr2-CLB,ΔCT-SmBiT (50 ng per well of a 90% confluent six-well plate). Twenty-four hours later, cells were lifted with TrpLE Express, resuspended in 37°C assay buffer (135 mM NaCl, 5 mM KCl, 0.4 mM MgCl2,1.8 mM CaCl2, 20 mM HEPES and 5 mM d-glucose, adjusted to pH 7.4), and transferred to a white flat bottom 96-well plate in triplicate with 20,000 cells per well. Coelenterazine-H (Thermo Scientific) in assay buffer prewarmed to 37°C was added to a final concentration of 5 μM and incubated for at least 5 minutes before data collection. For kinetic experiments, three time points were collected to establish baseline before vehicle or 10 μM isoproterenol addition (both of which contained 5 μM coelenterazine-H). Fold response was calculated by averaging the values across each triplicate and then dividing the isoproterenol treated samples by the corresponding buffer treated samples. For dose-response experiments, isoproterenol was added to the indicated final concentrations in a 37°C assay buffer and data was collected for 20 minutes. Fold response was calculated by averaging the values across each triplicate and then dividing the maximum by the minimum responses within each dose range. Dose-response curves were generated by a three-parameter nonlinear fit and tau values for time series data were determined using a one-phase association, both were calculated in Prism 9. At least three independent experiments were performed for all internalization assays.

### Live cell cAMP assay

β-arrestin-1/2 double knockout HEK293 cells were plated in 6-well plates, transfected with either mApple, βarr2-mApple, βarr2-ΔCT-mApple, βarr2-CLB-mApple, or βarr2-CLB,ΔCT-mApple. The following day, cells were lifted with TrypLE, transduced with CMV cADDis Green Upward cAMP sensor (Montana Molecular) according to manufacturer’s instructions, and transferred in triplicate at 50,000 cells per well in a black clear bottom 96-well plate (Corning). On the day of the experiment, the media was removed and replaced with 37°C assay buffer (135 mM NaCl, 5 mM KCl, 0.4 mM MgCl2,1.8 mM CaCl2, 20 mM HEPES and 5 mM d-glucose, adjusted to pH 7.4) and incubated for five minutes in the pre-warmed plate reader (H4 Synergy BioTek). Similar expression of mApple tagged plasmids was verified by fluorescence with monochromators set to Ex: 568/9.0 and Em: 592/13.5. Next, cADDis fluorescence baseline, and similarity of its expression across conditions, was established by three time points a minute apart using monochromators set to Ex: 500/9.0 and Em: 530/20.0. Isoproterenol was then added to a final concentration of 10 μM and cADDis fluorescence was measured every minute for the indicated time. Change in fluorescence was calculated by averaging across each triplicate and then dividing by the baseline. No fluorescence bleed through nor significant photobleaching were observed in separate experiments where cells expressing only mApple or cADDis green were measured with the same optical configuration. At least three independent experiments were performed.

### Sequence logos

PSI-BLAST was performed against bovine β-arrestin-2 protein sequence and iterated until convergence. Protein sequences annotated as β-arrestin and longer than 300 amino acids were kept while all others were discarded. Aligned regions of interest corresponding to 9 amino acids windows around either the C-lobe base, clathrin binding motif, or AP2β binding motifs were extracted and were input into a previously described tool (Crooks et al. 2004) to generate sequence logos.

### Statistical analysis

Quantitative data are expressed as the mean and error bars represent the standard error of the mean (s.e.m.) or standard deviation (s.d.) unless otherwise indicated. Scatter plots are overlaid with mean and s.e.m. Determination of statistical significance is described in each figure legend and was calculated using Prism 9.0 (GraphPad Software). *P < 0.05; **P < 0.01; ***P < 0.001 when compared with control, no-treatment, or other conditions. All experiments showing representative data were repeated at least three independent times with similar results. Independent experiments represent biological replicates.

## Data availability

All numerical data used to generate the figures has been included in the supporting data file. Source data for each figure panel is included as a separate worksheet in the combined excel document.

## Acknowledgements

We thank J. Benovic, M. Bouvier, A. Inoue, and S. Gutkind for sharing reagents and discussions; B. Kobilka, J. Janetzko, S. Sivaramakrishnan, A. Frost, R. Edwards, R. D. Mullins, M. Thompson, E. Blythe, as well as other von Zastrow laboratory and Manglik laboratory members for discussions. All live-cell imaging experiments were performed in the UCSF Center for Advanced Light Microscopy directed by D. Larsen. These studies were supported by grants from the U.S. National Institutes of Health R01DA010711 and R01DA012864 (M.v.Z.) and DP5OD023048 (A.M.). B.B.R. is a recipient of an American Heart Association Predoctoral Fellowship (19PRE34380570).

## Author Contributions

B.B.R., A.M., and M.v.Z. conceived of and designed the research. B.B.R. conducted all of the experiments and analyzed the results. B.B.R. and M.v.Z wrote the paper with input from A.M.

## Competing interests

The authors declare no competing interests.

**Correspondence and requests for materials should be addressed to M.v.Z**.

---

**Extended Data Fig. 1.**
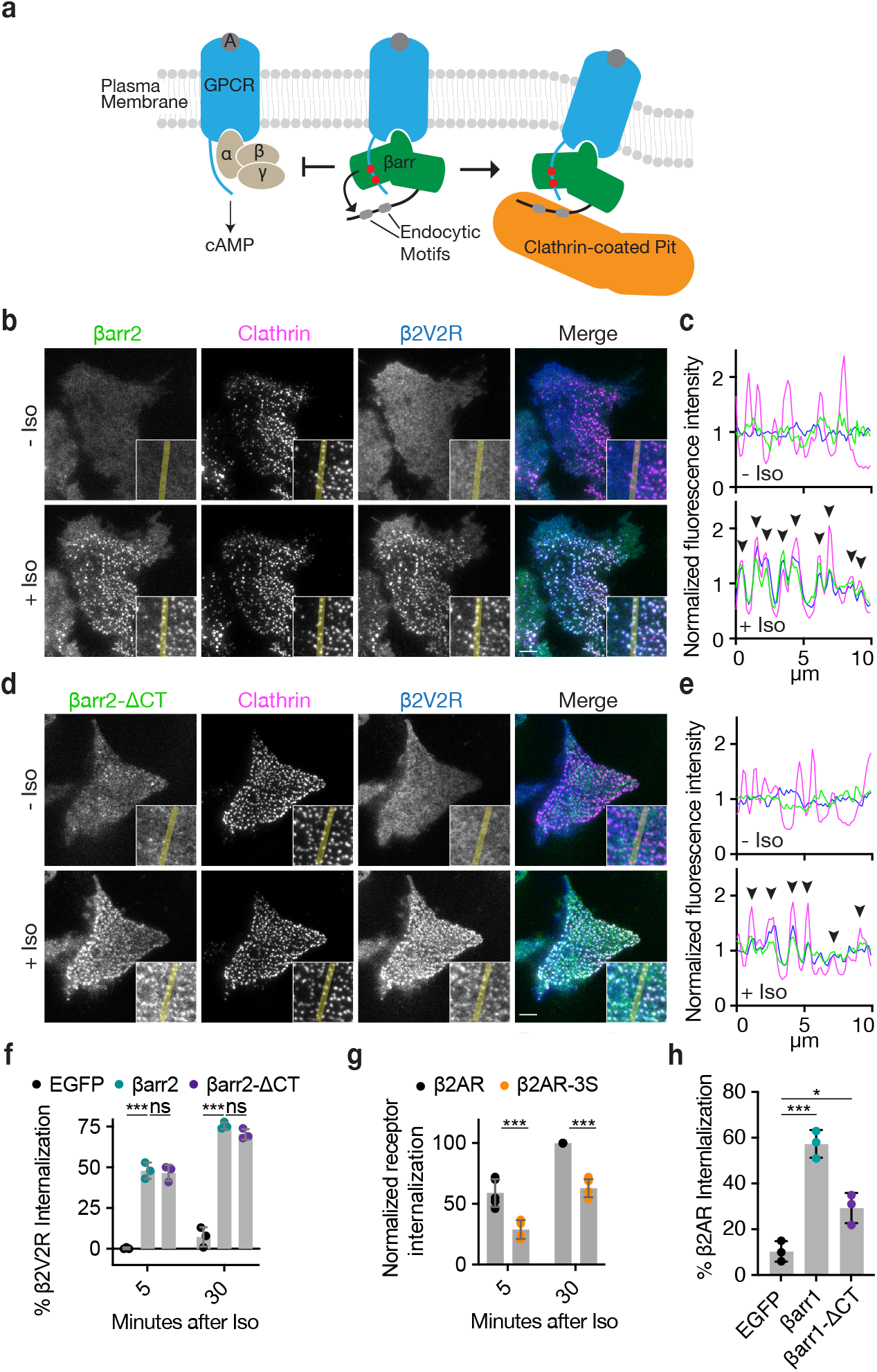
βarr2 and βarr1 CT are dispensable for GPCR internalization and β2AR phospho-sites are required for efficient internalization. **a**, Canonical model of β-arrestins’ mechanism of GPCR desensitization and endocytosis promoting functions. **b, d**, Representative live-cell TIRF microscopy images of βarr2-EGFP (green, b) or βarr2-ΔCT-EGFP (green, d) with FLAG-tagged β2V2R (blue) and clathrin-light-chain (magenta) pre- and post-stimulation with 10 μM isoproterenol (Iso) Scale bars represent 5 μm. Insets correspond to the central area of each cell. **c, e**, Normalized fluorescence intensity profiles from yellow lines shown in insets from panels (b) and (d) with colors corresponding to the image labels. **f**, Percent internalization of FLAG-tagged β2V2R co-expressed with either EGFP (black), βarr2-EGFP (green), or βarr2-ΔCT-EGFP (purple) after 5-or 30-minutes treatment with 10 μ? Iso. **g**, Normalized internalization of FLAG-tagged β2AR (black) or its phosphorylation site mutant, β2AR-3S (orange), after 5- or 30-minute treatments with 10 μM Iso coexpressed with β Arr2-EGFP. **h**, Percent internalization of FLAG-tagged β2AR after 30 minutes of stimulation with 10 μM Iso in DKO cells expressing either EGFP (black), βarr1-EGFP (green), or βarr1-ΔCT-EGFP. For **f, g**, and **h**, data is shown as mean ± standard deviation of n = 3 independent experiments with significance determined by either two-way ANOVA (df = 2, F = 15.3) or (df = 1, F = 63.82) with Dunnett’s test for multiple comparisons (f, g) or one-way ANOVA (df = 2, F = 49.81) with Tukey’s test for multiple comparisons (h), respectively. (ns P ≥ 0.05, * P < 0.05, *** P < 0.001). All experiments were performed in βarr1/2 DKO HEK293 cells.

**Extended Data Fig. 2.**
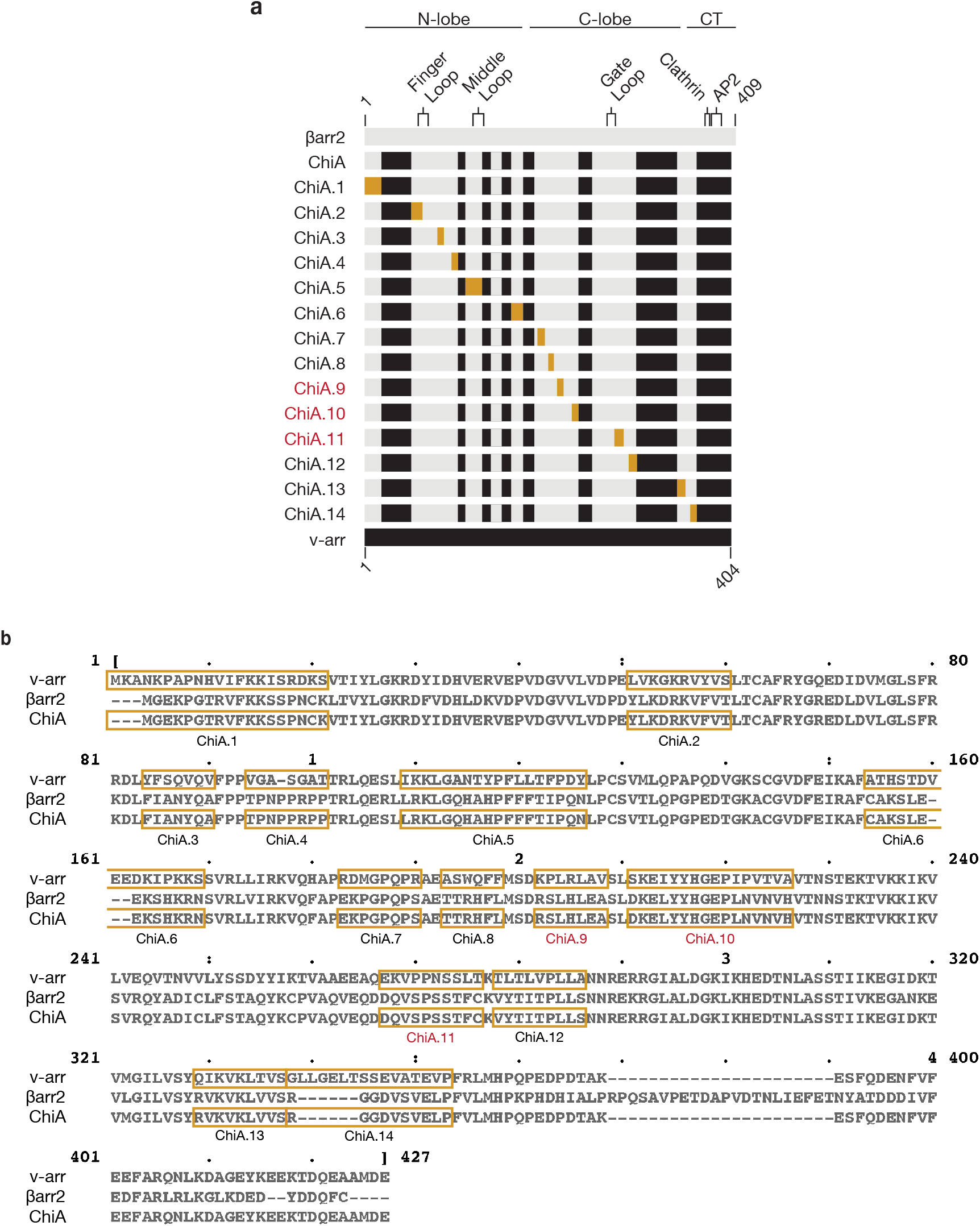
Diagram and sequences of visual arrestin / βarr2 chimeras. **a**, Diagram of βarr2 (gray) and visual arrestin (black) sequences. Visual arrestin sequences swapped into ChiA are gold to make ChiA.1-14. Major landmarks are shown for βarr2. **b**, Multiple sequence alignment of visual arrestin (v-arr), βarr2, and ChiA. Gold boxes in the v-arr sequence replace gold boxes in the ChiA sequence to make the indicated chimera. ChiA.9, 10, and 11 (red) abolished internalization of the β2V2R.

**Extended Data Fig. 3.**
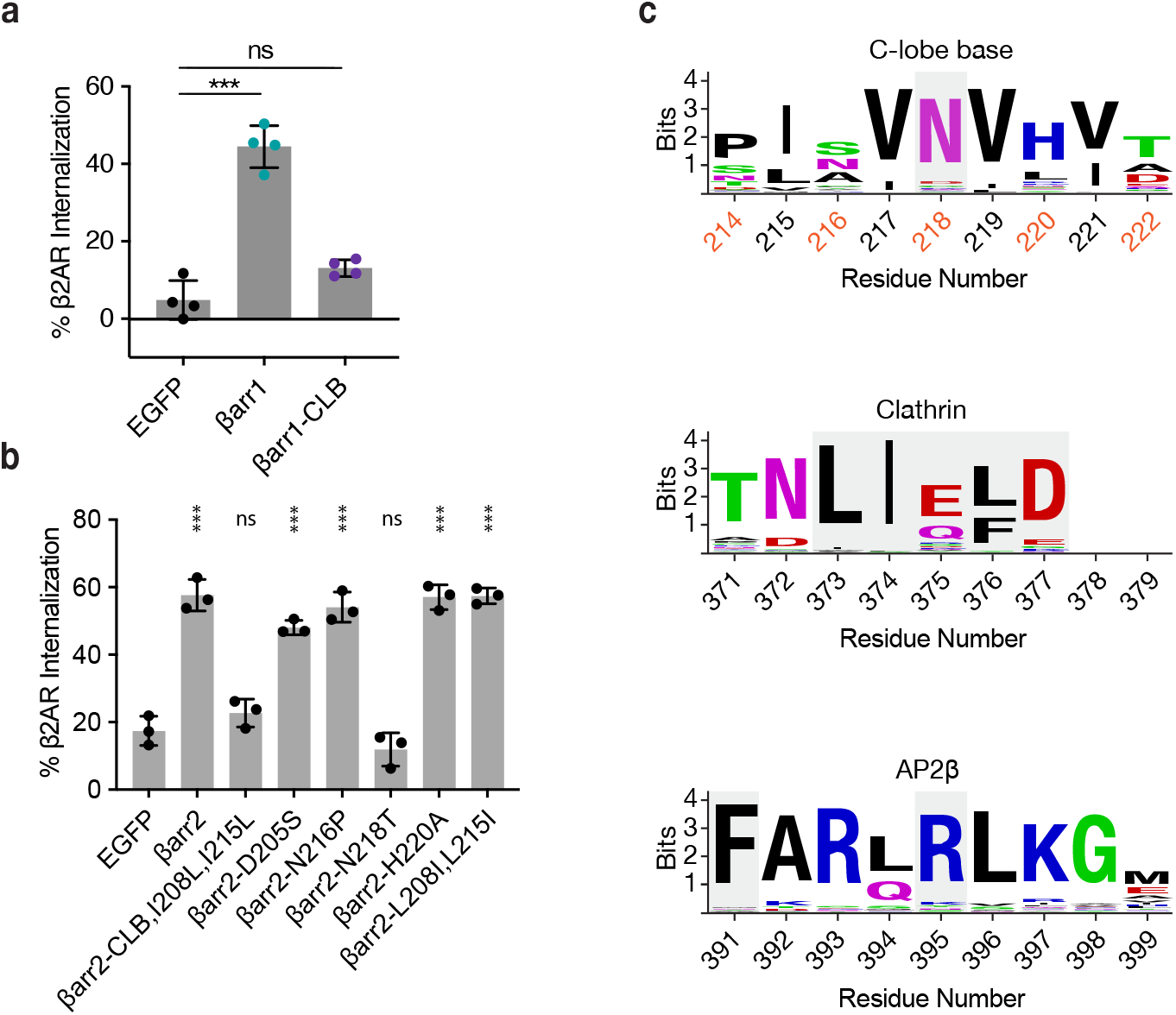
CLB is required for β2AR internalization in βarr1 and requires on a single conserved residue (N218) in βarr2. **a**, Internalization of β2AR after 30 minutes of stimulation with 10 μM isoproterenol when co-expressed with either EGFP, EGFP-tagged βarr1, or the βarr1 CLB mutant (D204S, S215P, N217T, and H219A) in βarr1/2 DKO HEK293s. Data shown as mean ± standard deviation of four independent experiments. Significance determined by ordinary one-way ANOVA (df = 2, F = 88.87) with Tukey’s multiple comparisons test. **b**, Internalization of FLAG-tagged β2AR co-expressed with the indicated construct after 30 minutes of stimulation with 10 μM isoproterenol. Data shown as mean ± standard deviation. Significance determined by ordinary one-way ANOVA (df = 8, F = 75.64) with Dunnett’s multiple comparison test against the negative control (EGFP) (n = 3). ns P ≥ 0.05, *** P < 0.001 **c**, Logos for sequence around N218 (gray) in the βarr2 CLB, clathrin binding box (gray), and AP2β binding site. Solvent exposed residues (CLB only) are numbered in orange. Amino acids that are critical to function or binding are colored in gray.

**Extended Data Fig. 4.**
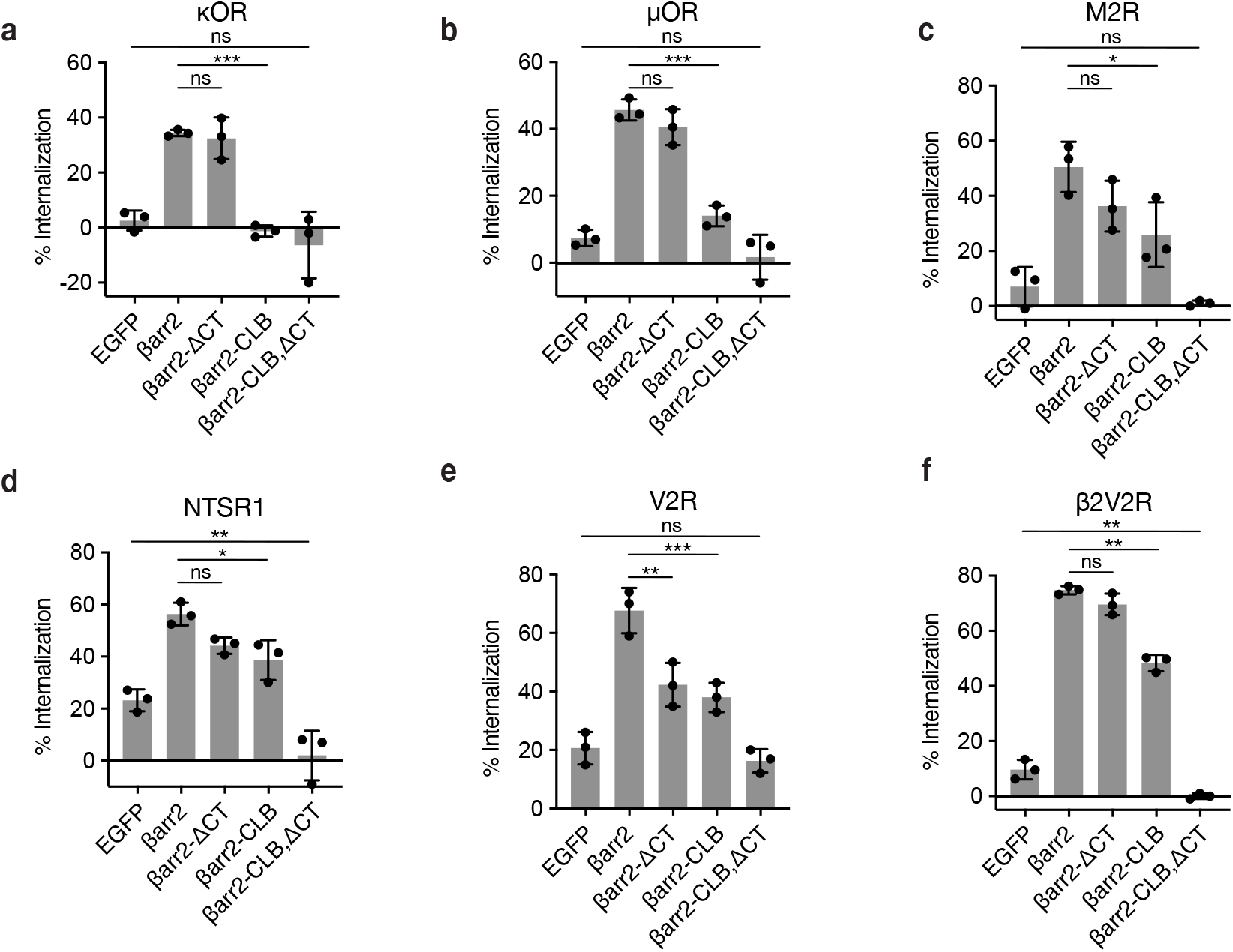
Internalization of GPCRs co-expressed with either wild type or mutants of βarr2. **a**-**f**, Percent internalization of the indicated receptor after 30 minutes of stimulation with either 10 μM dynorphin A-17 (a), 10 μM DAMGO 10(b), 10 μM carbachol (c), 10 μM neurotensin (d), 1 μM arginine vasopressin (e), or 10 μM isoproterenol (f) in βarr1/2 DKO HEK293 cells. Significance was determined by ordinary one-way ANOVA (df = 4 for all, F = 436.6, 33.13, 17.52, 33.88, 25.55, and 60.51, respectively) (ns P ≥ 0.05, * P < 0.05, *** P < 0.001). Data shown as mean ± standard deviation of three independent experiments.

**Extended Data Fig. 5.**
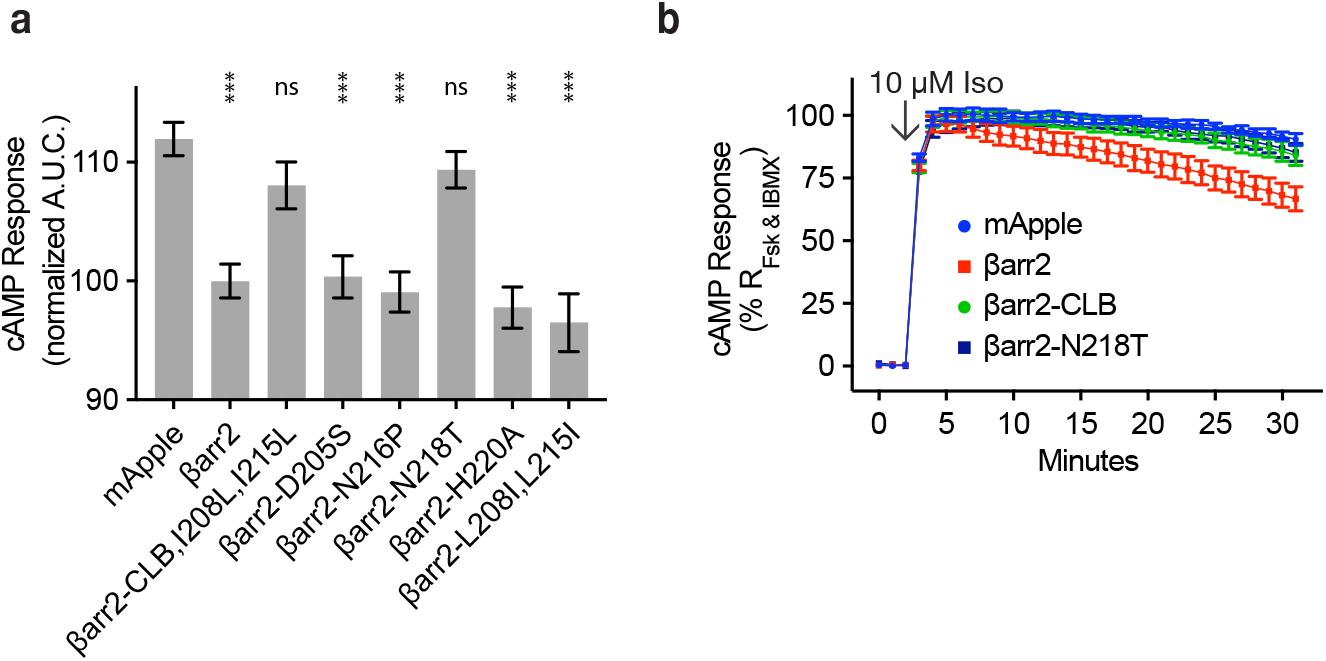
N218 in βarr2 is required for endogenous β2AR desensitization. **a**, Area under the curve (AUC) for the cAMP response elicited by endogenous β2AR after stimulation with 10 μM isoproterenol in βarr1/2 DKO HEK293s expressing the indicated construct and normalized to response in cells expressing wild type βarr2. Data shown as mean ± s.e.m. Significance determined by ordinary one-way ANOVA (df = 9, F = 13.47) with Dunnett’s multiple comparison test against the negative control (EGFP) **b**, Example kinetics of the cAMP response from endogenous β2AR after treatment with 10 μM Iso in βarr1/2 DKO HEK293 expressing the negative control (mApple, blue), βarr2-mApple (red), βarr2-CLB-mApple (green), and βarr2-N218T-mApple (dark blue). Data shown as mean ± standard deviation (n = 3). Kinetics for other mutants are not shown. All data shown is from three independent experiments.

**Supplementary Information Fig. 1.**
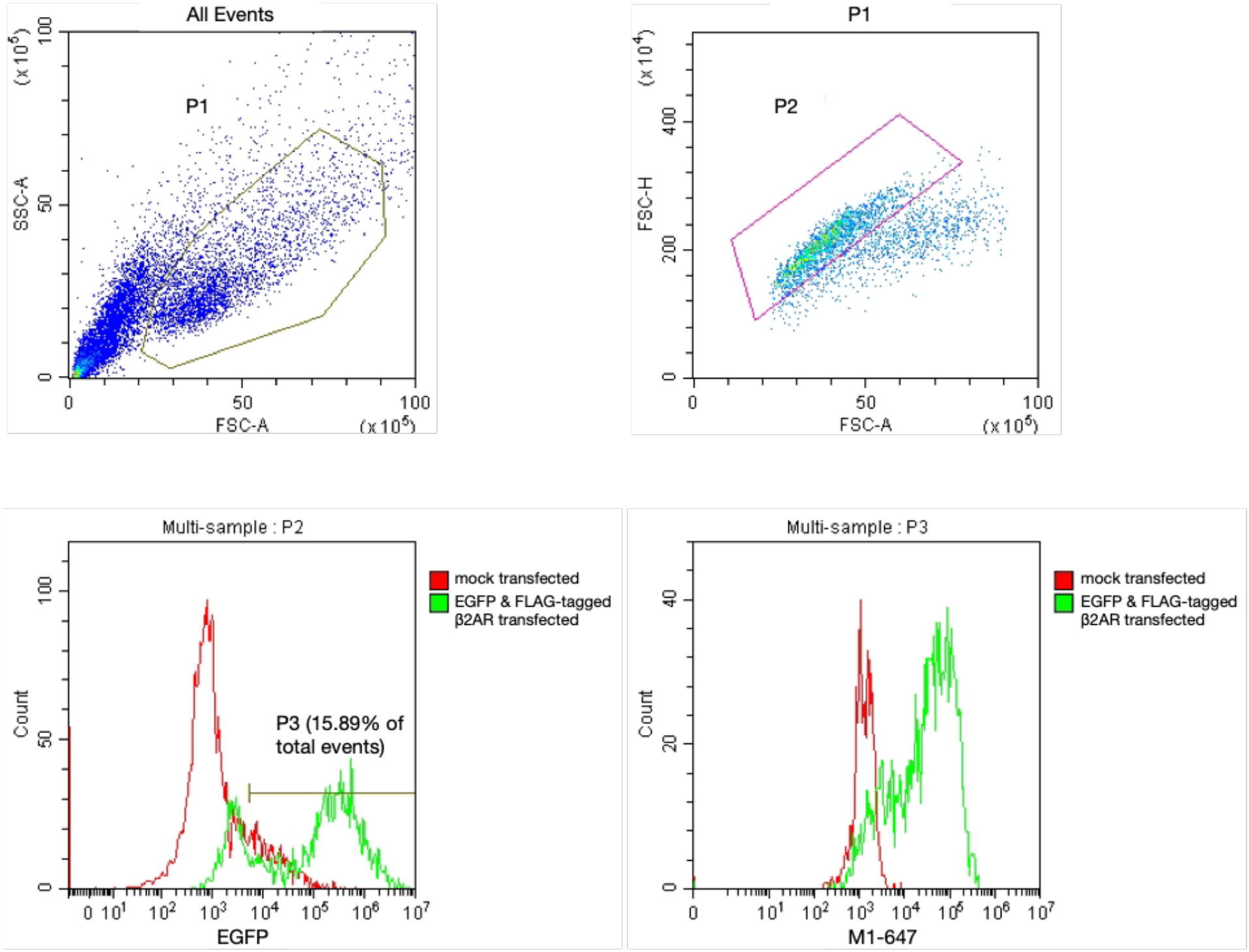
Gating strategy for flow cytometry internalization assays. P1 gate corresponds to cells. P2 gate corresponds to single cells. P3 gate corresponds to EGFP positive single cells. M1-647 fluorescence measurements were carried out on the P3 population.

